# Generalization of deep learning models for predicting spatial gene expression profiles using histology images: A breast cancer case study

**DOI:** 10.1101/2023.09.20.558624

**Authors:** Yuanhao Jiang, Jacky Xie, Xiao Tan, Nan Ye, Quan Nguyen

## Abstract

Spatial transcriptomics is a breakthrough technology that enables spatially-resolved measurement of molecular profiles in tissues, opening the opportunity for integrated analyses of morphology and transcriptional profiles through paired imaging and gene expression data. However, the high cost of generating data has limited its widespread adoption. Predicting gene expression profiles from histology images only can be an effective and cost-efficient *in-silico spatial transcriptomics* solution but is computationally challenging and current methods are limited in model performance. To advance research in this emerging and important field, this study makes the following contributions. We first provide a systematic review of deep learning methods for predicting gene expression profiles from histology images, highlighting similarities and differences in algorithm, model architecture, and data processing pipelines. Second, we performed extensive experiments to evaluate the generalization performance of the reviewed methods on several spatial transcriptomics datasets for breast cancer, where the datasets are generated using different technologies. Lastly, we propose several ideas for model improvement and empirically investigate their effectiveness. Our results shed insight on key features in a neural network model that either improve or not the performance of *in-silico spatial transcriptomics*, and we highlight challenges in developing algorithms with strong generalization performance.

**Key Messages:**

- We comprehensively compared the performance of existing methods for predicting spatial gene expression profiles from histology images
- We assessed the roles of different algorithms, model architectures, and data processing pipelines to model performance
- We performed extensive experiments to evaluate the generalization of the models on in-distribution and out-of-distribution spatial transcriptomics datasets
- We proposed several strategies for improving existing models and empirically investigated their effectiveness

## Introduction

Spatial transcriptomics (ST) is a rapidly developing technology for producing histopathological images paired with gene expression profiles of thousands of individual spots/cells, providing information on both the unseen molecular signatures and imaging morphological features [1]. Technology such as the 10x Genomics Visium platform [2] measures the gene expression on spots of resolutions 55 to 100 micrometers in diameter that are distributed close together at fixed locations across the tissue, which can then be mapped back to the underlying high-resolution stained tissue image. ST has been used to characterize the spatial heterogeneity of cancers, providing new insights in cancer research, and opening an unprecedented potential for novel capabilities in the diagnosis and prognosis of cancer (e.g., see the survey paper [3]). However, the wide adoption of ST has been limited by its high cost.

Several deep learning methods have been developed for predicting gene expression profiles from Hematoxylin and Eosin (H&E) stained histology images. These include methods that predict bulk or single-cell RNA profiles which do not provide spatial information [4, 5], and methods that are able to predict spatially resolved profiles [6, 7, 8, 9, 10, 11]. In this work, we focus on the latter class of methods that are emerging with the potential to bring about new abilities to predict spatial transcriptomic data for histopathological image assessments. While the published results support the potential of the deep learning approach, the performance of these existing methods still requires significant improvement. At the same time, there are several limitations in existing studies that prevent a good understanding of the state of the field: (a) existing studies often use different datasets to compare a subset of the algorithms, instead of comparing existing algorithms on the same datasets; (b) existing studies focus on in-distribution (ID) generalization performance (where the training and test sets are sampled from the same distribution), while in practice, it is even more important to assess the generalization for out-of-distribution (OOD) (where the training and test sets may follow different distributions); (c) the methods often differ in multiple aspects, including the model architectures, preprocessing techniques, and data augmentation strategies, but it is often unclear which of the differences account for the performance differences.

Our paper aims to address the above limitations and advance research in the field by performing a comprehensive assessment of the generalization performance of existing approaches and experimenting some new ideas for improving models. Specifically, we identify six existing methods and we study the following questions:

- How do the methods compare with each other on datasets generated by different technologies?
- How well do the methods generalize on OOD data?
- How effective are the different preprocessing and data augmentation techniques?
- How effective are some new ideas for improving deep learning models (e.g., the use of pre-trained models)?

We focus on breast cancer in this study, because of the availability of relatively large datasets generated by a legacy spatial transcriptomics protocol (100 μm resolution spots) and a recent version of 10x Genomics Visium protocol (55 μm spots), which have different resolutions and detection sensitivities, and would allow us to study the above questions. In fact, some previous work has partially compared different methods using breast cancer tissues [12, 7, 8, 10], but the important questions and limitations as mentioned above remained unaddressed. For example, several methods [12, 7, 8] have not been quantitatively evaluated on datasets generated by newer, higher-resolution technologies, and their performance on OOD data has not been evaluated.

Our paper contributes a systematic review of existing methods, and results in several interesting empirical insights, with some highlighted below.

- Top-performing methods: Hist2ST, BLEEP, and STimage consistently achieved higher test set performance for both the in-distribution and OOD settings.
- General performance: All the evaluated methods predicted overall variable genes or cancer markers with relatively low accuracy, resulting in low-performance metrics across the diverse datasets. This raises the need for assessing which genes can be predicted and at which performance ranges.
- Limited generalization: Most methods exhibited limited generalization capability, with low performance when applied to OOD data. Some methods exhibit overfitting to the data, and we highlight differences in models that may account for improved robustness.
- Negative transfer: Using pre-trained image encoders that have been trained on general images led to negative transfer when applied to H&E images, resulting in lower performance compared to training from the ground up.
- Impact of image augmentation and expression preprocessing: Various image augmentation strategies did not have a discernible impact on overall performance. In addition, different preprocessing transformations for gene expression values did not lead to significantly different results.

The remainder of this paper is structured as follows. First, we review and compare six existing deep-learning algorithms for predicting spatial gene expression profiles from histology images. We then describe our benchmark methodology, with details on the datasets used, the performance metrics, and variants of a top-performing algorithm Hist2ST [8]. This is followed by assessing the ID and OOD generalization performances of the existing methods and the variants of Hist2ST. We then discuss the benchmarking results and critically review the differences between methods and model architectures in light of the observed results. We also describe experimental results from testing several new ideas for improving the performance. Finally, we address the limitations of this benchmarking work and highlight the potential challenges of applying deep learning models for the prediction of spatial transcriptomic profiles.

## Review of existing methods

We identify and review six existing methods in this section. All methods extract image patches from the whole-slide histology images based on the spatial coordinates of the measured gene expression. The transformer-based methods (HisToGene, Hist2ST) treat all spots on a single tissue section as an independent training sample, making use of the global relationship between spots; the other methods treat all spots (both within and between tissue samples) as independent training samples, with the number of input images determined by the batch size. Most methods (ST-Net, HisToGene, Hist2ST, STimage, DeepSpaCE) learn to predict gene expression directly by multivariate regression; the exception is BLEEP, which infers the gene expression by querying through nearest-neighbors in a learned joint embedding space for image and gene expression. The architectures used in the first five methods all share the same high-level structure: an image encoder is used to extract features from the image patches, then a prediction head is used to predict the gene expression profiles using the extracted features. Some methods additionally include a distribution module for predicting the distribution of the gene expression values rather than a fixed value. This general structure is illustrated in the top figure in Figure 1 (a). The five methods differ in the specific networks used for the encoder, the prediction head, and the distribution module, as will be discussed below. In addition to differences in model architecture, these methods employ different techniques for processing image and gene expression data; summarized in Table S1. The methods we have identified have publicly available code and have been individually evaluated previously on different datasets, including breast cancer data; we provide a summary in Supplementary Table S4. In this section, we focus on comparing the model architectures and loss functions used and review each method separately below.

**Fig. 1.**
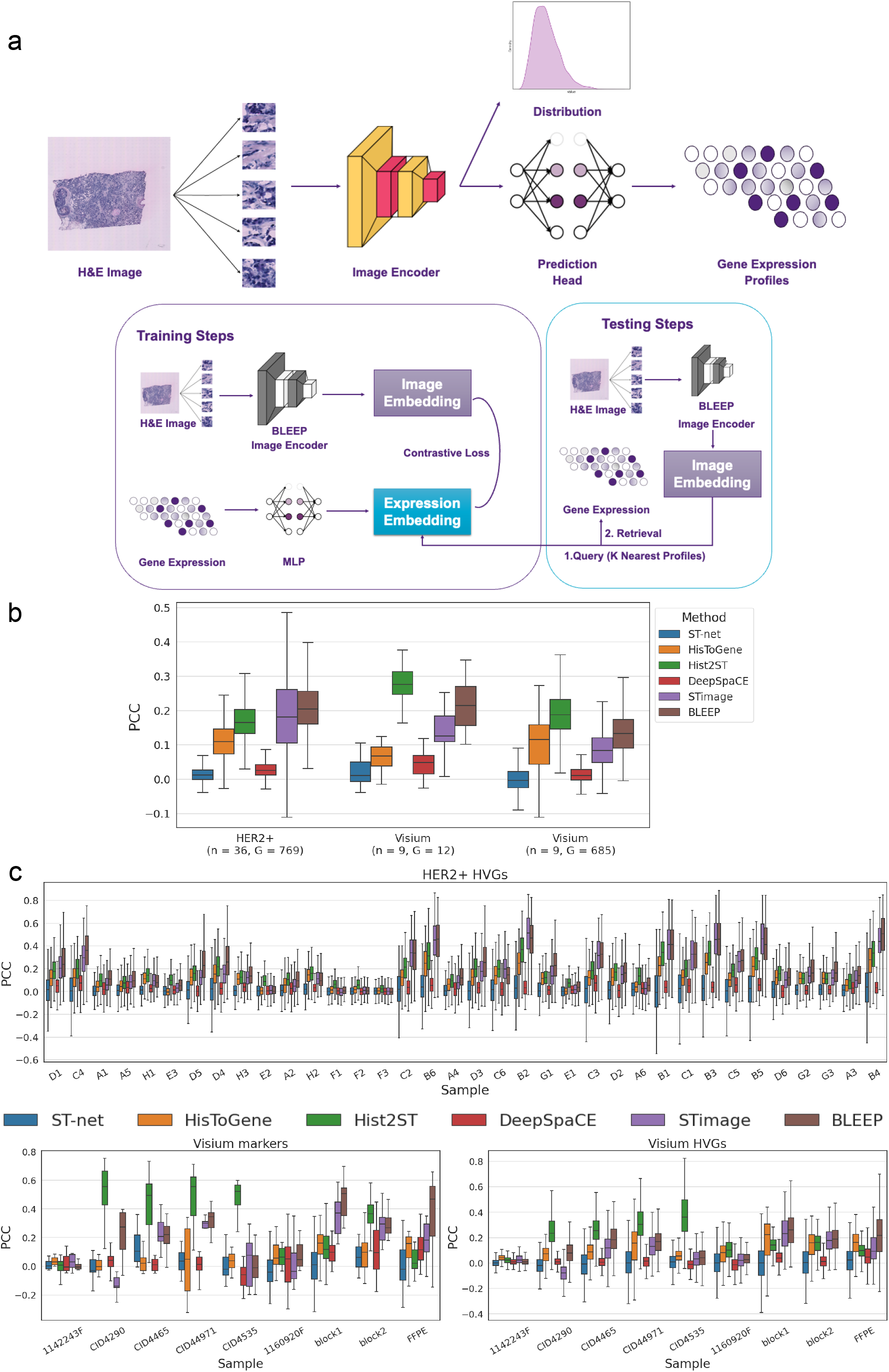
a) Schematic overview of the general model architecture shared by the six existing methods that predict gene expression from H&E images: ST-Net, HisToGene, Hist2ST, STimage, DeepSpaCE and BLEEP. The top diagram illustrates the common feed-forward architecture of regression-based methods, and the bottom displays the query-reference method BLEEP. **b)** Model performances on the Visium breast cancer and HER2+ datasets, showing the averaged LOOCV results. Box plots indicate the Pearson correlation coefficient (PCC) for highly variable genes (leftmost and rightmost), and marker genes (middle). **c)** Individual results of the LOOCV for each hold-out test sample. The top panel corresponds to HVGs for the HER2+ dataset; the bottom left and right, marker genes and HVGs for the Visium dataset, respectively.

ST-Net [6] is a CNN model that employs a DenseNet-121 pre-trained on ImageNet [13] as an image encoder. The whole-slide histology images are divided into 224×224-pixels patches for each spot, which are transformed into features by the image encoder. The prediction head is a single fully connected output layer, which directly predicts the vector of gene expression. ST-Net is trained by minimizing the mean squared error (MSE).

HisToGene [7] utilizes a modified vision transformer (ViT) model as an image encoder, which is able to incorporate the spatial relation of image spots. 112×112-pixel image patches are flattened and passed into a linear layer to produce the patch embeddings of dimension 1024. In addition, a learnable linear layer is used to map the spatial coordinates of each spot into a positional embedding vector of the same dimension. The patch embeddings and position embeddings are aggregated via summation and then input into eight multi-headed attention layers to generate latent embeddings. The prediction head is a single feed-forward layer that directly outputs the gene expression values. The model is trained to minimize the MSE. Hist2ST [8] consists of three main modules – Convmixer, Transformer, and Graph Convolutional Network (GCN) – to learn a global feature representation for all image spots on a single tissue section. 112×112-pixel patches are extracted around each spot. The initial image encoder is a Convmixer, a variant of CNN, that produces image features for each spot. The transformer module encodes spatial locations and fuses them with the image features, employing eight multi-head attention layers to learn global spatial dependencies. The aggregated features from the transformer module are then input into the GCN module to explicitly learn local spatial dependencies. The prediction head is a fully connected layer. In addition, there is a distribution module that estimates parameters for Zero-Inflated Negative Binomial (ZINB) [14] distributions for each gene; this is learned by minimizing the negative log-likelihood. Self-distillation [15] is used to learn from augmented image patches. The loss function combines the MSE from the prediction head, the ZINB loss, and the self-distillation loss.

STimage [10] is a CNN model that uses a ResNet50 image encoder to extract features from 299×229-pixel image patches. The features are input into a distribution module, which consists of two fully connected output layers to estimate the parameters of a Negative Binomial (NB) distribution for each gene. This is achieved by minimizing the negative log-likelihood. The estimated mean of the NB distribution for each gene is used as the prediction, and the parameters can be used to quantify the uncertainty in the prediction. STimage additionally employs an ensemble approach to account for variation across independent training runs to improve performance and robustness.

DeepSpaCE [9] is a CNN model that utilizes the VGG16 architecture as an image encoder. Image patches of size 150% times the original spot dimensions are extracted from the histology image. The prediction head consists of two fully connected layers for predicting the gene expressions. The loss function used is the smooth L1 loss.

BLEEP [11] learns a bimodal embedding for image and gene expression data. Its architecture is illustrated by the bottom figure in Figure 1 (a). The process involves extracting features from the image tiles and expression vectors using separate encoders. The image encoder is a pre-trained ResNet50, while a fully connected network serves as the expression encoder. The image and gene expression features are then separately projected into image embeddings and expression embeddings, with a shared dimension of 256. Contrastive learning [16] is employed to align the latent space for the two embeddings by minimizing the cross entropy. During inference, the image patches are mapped to embeddings by the encoder, and the k-nearest expression profiles from the reference dataset are selected based on their proximity, measured by Euclidean distance, to each patch in the joint embedding space. The prediction of expression profiles for the query patches is then performed by taking a linear combination of the selected expression profiles.

## Methods

We evaluate the ID and OOD generalization performance of all six models and multiple variants of Hist2ST using two data settings, as detailed below.

### Datasets

In this study, we utilized two spatial transcriptomic breast cancer datasets generated by the Visium technology and the legacy spatial transcriptomics technology. The two datasets, summarized in Table 1 and described in detail below, differ in terms of the technology, the number of spots, the spot resolution, the number of genes detected per spot, and the number of patients per datatype. Patient samples were collected in different cohorts and data was generated by different laboratories, representing technical variations likely observed in the real application settings. This approach allowed us to assess how various models perform across datasets with different characteristics.

**Table 1.** Summary of samples from the HER2+ Legacy and Visium datasets.

| Dataset | Patients | Tissues | Cohorts | Resolution | Spots | Genes |
| --- | --- | --- | --- | --- | --- | --- |
| HER2+ | 36 | Frozen | He et al. | 100 μm | 13,620 | 11,871 |
| Visium | 6 | Frozen | Wu et al. | 55 μm | 16,456 | 36,601 |
| Visium | 2 | Frozen | 10x | 55 μm | 7,785 | 36,601 |
| Visium | 1 | FFPE | 10x | 55 μm | 2,518 | 36,601 |

The lower resolution (100 μm per spot) HER2+ breast tumour ST dataset [17] comprises 36 tissue sections from 36 HER2+ patients, with data generated using the initial version of the spatial transcriptomics protocol before the method was further developed by 10x Genomics to increase the resolution to 55 μm per spot. The term legacy ST is referred to for historical reasons, without implying about quality of the data. Among the 36 samples, there are annotated cell type labels for 8 tissues, by trained pathologists. Each sample consists of around 300 to 600 spots, with each spot containing the measurements for 11,871 genes. The images are low resolution, less than 9000 pixels in height and width.

The higher resolution Visium breast cancer dataset, introduced and described in [10], consists of nine breast cancer tissue samples, which includes 3 samples (2 fresh-frozen and 1 formalin-fixed-paraffin-embedded, FFPE, tissue) from 10x Genomics [18] and six samples obtained from Swarbrick’s laboratory [19]. Each sample is measured on the 10x Visium platform and contains around 1300 to 4900 spots, with each spot containing the measurements for up to 36,601 genes. The images are high resolution, ranging from approximately 9000 to 40000 pixels in height and width.

### Genes selected for evaluation

Although spatial transcriptomic data contains measurements for thousands of genes, the methods considered have not been designed to scale to predict more than a small subset of usually 200 to 1000 genes. As the performance is likely dependent on the gene targets for prediction, we considered the prediction of two sets of genes: highly variable genes (HVGs), and a smaller set of 12 cancer-associated genes (marker genes). For the HER2+ dataset, the performance was assessed on the prediction of the 769 shared HVGs and the Visium dataset was assessed on both the HVGs (685 shared), and the 12 marker genes. Of note is that the selection of HVGs for prediction is a common practice across all methods, although the rationale for why the HVGs are suitable or not is not clearly justified, except for the fact that these methods are not scalable to predict all genes.

For the BLEEP method, the model appears to be more scalable and the selected gene panel may incorporate a larger set of highly variable genes for both training and inference. Therefore, we also assess the model performance when a larger set of genes are used, and so we have benchmarked the BLEEP method in two ways for comparison: for the HVGs, the gene set was fixed which is the same case as the other methods; and another gene set was union of the 12 marker genes and 1000 HVGs across all samples, resulting in an enlarged set.

### Performance measures

We describe how we evaluate a method’s ID and OOD generalization performance below. We measure the predictive performance of a method for a gene using the Pearson correlation coefficient (PCC), defined by

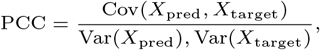

where *X*_pred_, *X*_target_ is the predicted and measured target gene expression for all spots in the sample, respectively. PCC is a better measure than the mean squared error or the mean absolute error, because it is not dependent on the differences in the absolute scale between abundant and lowly-expressed genes. The ID generalization performance is measured using LOOCV (leave-one-out cross-validation) as the number of samples is small in the datasets used. Specifically, we leave one sample out at a time and train a model on the remaining samples. We then make predictions on the hold-out sample and measure the PCC for each gene.

The OOD generalization performance is measured only on the Visium dataset, using the marker genes. Specifically, we partitioned the Visium dataset (illustrated in 2a) into a training set, a validation set, and a test set. The training set and the validation set contain 4 samples and 2 samples respectively from the Swarbrick data [19]. The test set contains samples from 10x Genomics. We used the validation set for measuring the ID generalization performance instead of for hyperparameter tuning. The test set is used for measuring the OOD generalization performance. The partition was motivated by the observation that data produced from two different laboratories are likely to have distribution shifts in the image and gene expression data. To verify this, Figure 2a shows that the whole slide images for the training and validation dataset are separated into two groups with different color profiles, which are further different from the test set. Figure 2b verifies that the color distribution of the RGB channels is distinctive and varies across individual samples and laboratories. In terms of gene expression, Figure 2c shows that the distribution of gene expression has similar distinctiveness between the training and validation sets and the test set samples.

**Fig. 2.**
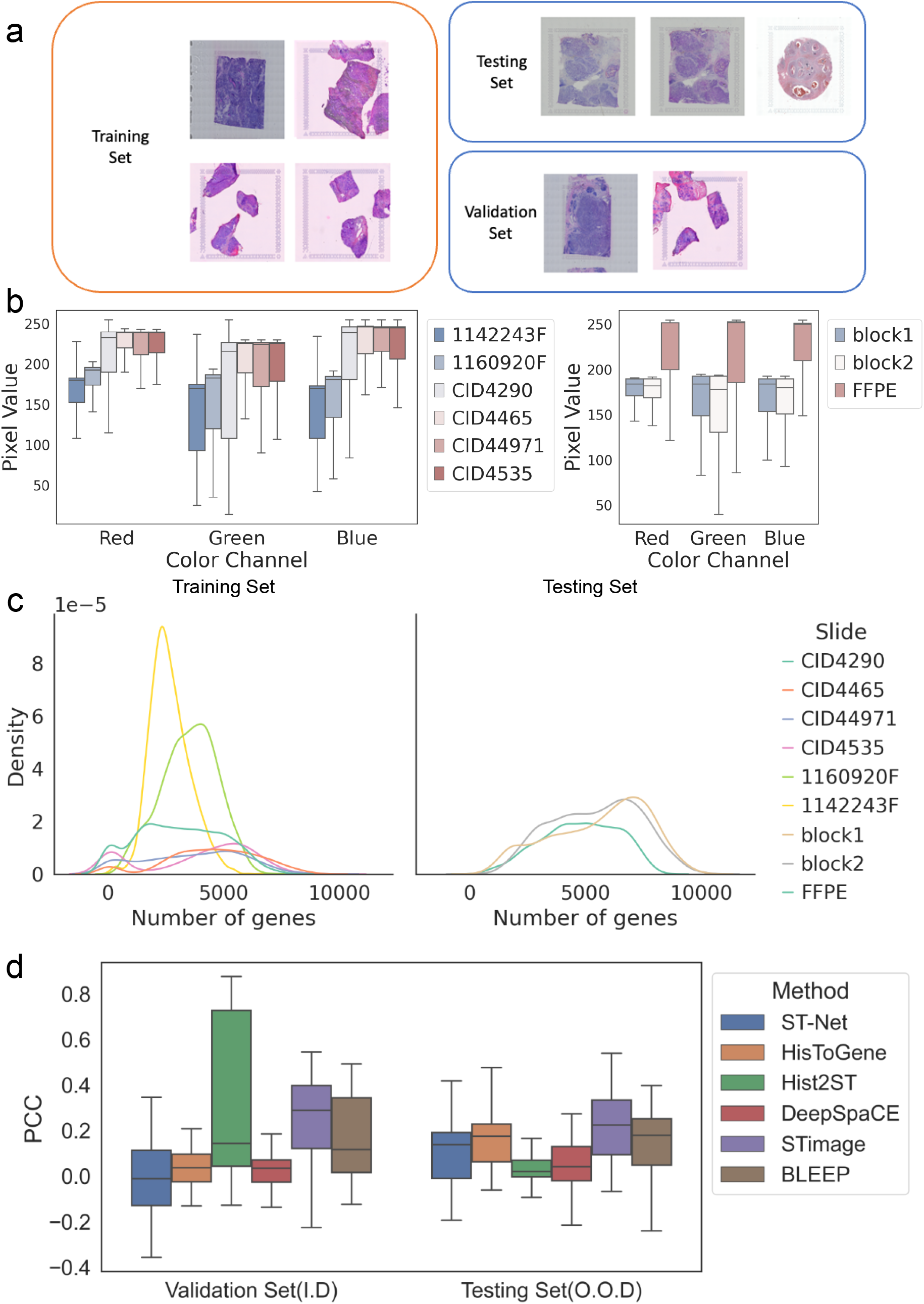
a) Sample images in the Visium dataset and the dataset splits for assessing the generalization of methods. **b)** Color distributions in the Visium dataset, where the training set refers to the combined training and validation sets. **c)** Distributions of gene expression in the Visium dataset, showing the density (y-axis, frequency of the spots) corresponding to the number of genes detected per spot (x-axis). **d)** Assessment of generalization ability on in-distribution validation set and out-of-distribution (OOD) test set. Box plots show the correlation between predicted and true gene expression for each gene predicted (marker genes).

### Algorithm settings

All the methods were first trained and tested with their default hyperparameters, pre-and post-processing steps, and data augmentation methods. We provide a concise overview of the processing steps applied to both gene expression data and image data for each method in Table S1 and a detailed summary in Supplementary section 7.1. Each model was trained on a single NVIDIA Tesla V100 SMX2 GPU with 32 GB RAM.

When testing Hist2ST and HisToGene with the Visium high-resolution data, we faced out-of-memory issues. Since these methods include transformer modules that process all the spots in tissue samples at once, processing Visium samples that usually contain more than 2000 spots exceeds the available memory. To address this, we divided each whole image into smaller, non-overlapping square windows of 4000×4000-pixel sections, resulting in multiple training instances for each sample, and each instance can be trained in memory successfully, with each batch containing hundreds of spots rather than thousands.

### Proposed modifications for Hist2ST

To explore possible approaches to improve predictive performance and robustness, we adopted Hist2ST as a baseline model and experimented with the following modifications, which are illustrated in the Supplementary section 7.1. We also empirically compare different augmentation techniques and transformations for gene preprocessing to determine if there is an augmentation and preprocessing method that can optimally improve the model performance.

### Image augmentation and color normalization

Existing methods employ different data augmentation methods to improve model robustness to variation in image data and improve generalization. These techniques are summarized in Figure S2a. To investigate whether different augmentation techniques result in better performance, we benchmarked each technique using the Hist2ST backbone model.

H&E images produced by different equipment and laboratories can vary greatly in stain levels and color distribution. We observe such variations in the Visium dataset (Figure 2b). Although some existing techniques apply random color augmentation such as color jitter (see Table S1), many do not employ processing steps that take into consideration the unique characteristics of H&E images, which possess a color distribution greatly distinct from natural images. For example, only STimage applies stain normalization to the images prior to training. Thus, we also systematically benchmarked the effects of applying dedicated stain color augmentation and normalization methods on model performance and robustness. This included stainlib [20], which offers H&E-intensity color augmentation and Reinhard color normalization, and RandStainNA [21], which integrates stain normalization and stain augmentation in a combined fashion to constrain the variability of stain styles within a practicable range.

### Preprocessing for gene expression

Different methods utilize different transformations for the gene expression values; this is summarized in Table S1. Here, log-transformation refers to a log-transformation on the counts with a pseudocount addition. To determine whether there is a transformation that results in better predictive performance, we benchmarked the effect of the following transformations: no transformation (raw), log-transformation, log-transformation on normalized counts (log counts per million), and MinMax scaling (counts scaled to the range [0, 1]).

### Auxiliary classification loss

We investigated whether the addition of an auxiliary classification task in training could improve the performance of gene expression prediction. As a subset of the Visium and HER2+ datasets contain spot annotations by pathologists for the same tissue section with the spatial data, employing this information by introducing a classifier module during training could potentially lend the model to learn more informative features for each image spot. Furthermore, the classification loss could act as a regularization term to improve the generalization of the model to images of similar tissue characteristics in new data. Here, we proposed Regclass, a variant of Hist2ST, which introduces an auxiliary classification head in the model and is trained to jointly optimize the original loss for regression and the cross-entropy loss for classification.

The workflow of Regclass is shown in Algorithm 1. The classification head is a 4-layer multilayer perceptron, and the remaining components are the same as in Hist2ST, including the Convmixer, position encoder, transformer, graph convolution network, and regression head.

#### Algorithm 1 Regclass model

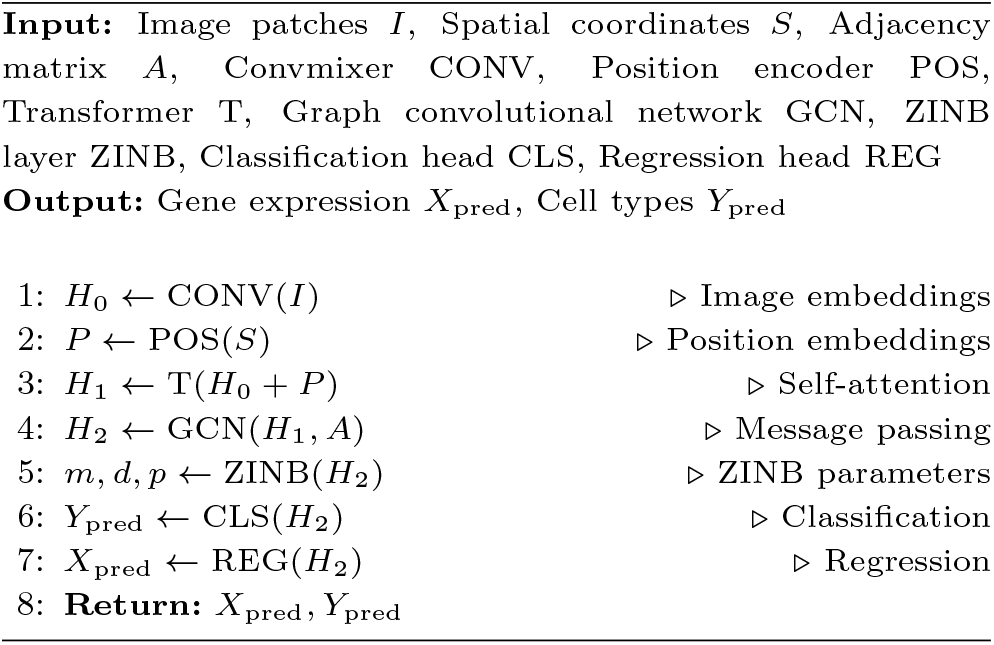

The loss is calculated as follows.

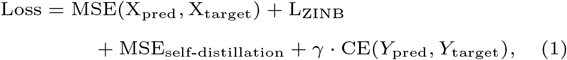

where *X*_pred_, *X*_target_ are the predicted and ground truth gene expression values, respectively; *Y*_pred_, *Y*_target_ are the predicted and ground truth tissue classes, respectively; and *γ* is a hyperparameter to scale the cross-entropy loss CE. The MSE, L_ZINB_, and MSE_self-distillation_ losses are the same as in Hist2ST. We investigated different values of *γ* to determine an optimal value for both accuracy and generalization.

### Pre-trained image encoders

Here, we investigated whether using transfer learning with different image encoders trained on ImageNet could improve the performance of the baseline Hist2ST model. As the datasets are small, using pre-trained image encoders could be advantageous as it allows the model to leverage the knowledge gained from broader training data, enabling it to extract better features from a model initialized with meaningful weights.

We benchmarked the original Hist2ST model, which used a Convmixer as the initial image encoder, with the following models pre-trained on ImageNet: ResNet50 [12], EfficientNet V2 [22] and Swin Transformer [23]. Only the last layer of each backbone was fine-tuned.

### Simplification with graph attention network

The original Hist2ST model includes a transformer and GCN module to learn spatially aware representations. However, we have found that the transformer does not scale well to newer ST data, such as for the Visium samples, which contain thousands of spots on one tissue section. This is due to the quadratic complexity of the transformer in terms of the input sequence length from global self-attention.

On the other hand, the graph attention network (GAT) uses the attention mechanism to learn node embeddings by considering only the neighbourhood information of each node. Attending to only a subset of neighbouring nodes for each node reduces the overall computation when compared to the transformer. We hypothesized that a simpler model, which consists of replacing the transformer and GCN modules in Hist2ST with a 4-layer GAT, could perform similarly to the original. This was modified in conjunction with the replacement of the initial feature extractor and benchmarked for comparison.

## Results

### Performance of existing methods

#### ID generalization performance

Figures 1b,c shows the results of the performance comparison between the six methods across the two datasets. In general, the best-performing methods are STimage, BLEEP, and Hist2ST, which consistently outperform other methods based on average PCC values across all tissues from different datasets. HisToGene had a slightly lower performance on average, with a median accuracy of around 0.1 correlation. In contrast, ST-Net and DeepSpaCE were unable to consistently predict HVGs and markers with median accuracy above 0.1 correlation. The relative performance ranking between methods is similar across HER2+ and Visium datasets, suggesting that the performance of methods remains consistent and does not change much with an increase in image resolution and number of spots. On the HER2+ dataset, the performance of BLEEP and STimage is significantly better than that of the other models. On the Visium dataset, the performance of Hist2ST consistently outperforms other models. We find that the ranking between methods is consistent across predicting the smaller set of marker genes and the larger set of HVGs on the Visium dataset, with slightly higher average PCC on the markers.

### OOD generalization performance

Figure 2d shows that in general, the average PCC on the validation (ID samples) and test sets (OOD) are similar. STimage has the highest median performance on the OOD test data. For Hist2ST, there is a large discrepancy between the high performance on the validation dataset and the lower performance on the test dataset, suggesting possible overfitting to the training data and less generalizability.

### Performance of Hist2ST modifications

We empirically assessed the performance changes due to each of the new modifications in model architecture, regularization, and data augmentation to the backbone Hist2ST model. We applied the data preprocessing methods as shown in Figure S1a to the images in the training set and the trained models were tested on the unseen, unprocessed image data. The Hist2ST’s performance results were compared between different augmentation techniques and gene expression transformation methods. Our comparisons suggest that the data augmentation techniques resulted in similar performance (Figure S1c). Comparing stain augmentation and color normalization we also observed that both steps did not improve the performance (Figure S1d). In addition, we found that the various transformations for the gene expression values resulted in similar performance in terms of PCC, where the only significant difference is that the variance for the log-normalized transformation is smaller than the others (Figure S2e).

To assess the usefulness of the auxiliary classification module in a modified Regclass model as illustrated in Figure S2, we found that the model can be trained to classify tissue types successfully in most of the samples, with both the F1 score and Area under the ROC Curve (AUC) generally above 0.8 Figure S3. However, this did not translate to performance improvements for predicting gene expression, with the performance being similar to the original Hist2ST (Figure S3a). Figure S3b shows that Regclass slightly improves the performance on OOD data, for values of *γ* above 0.

Comparing the performance of the pre-trained feature extractors as shown in Figure S3a, we found that alternative extractors show a large performance reduction compared to the method used in Hist2ST. Similarly, the use of the graph attention network did not result in any improvements in performance on both datasets.

## Discussion

We benchmarked six methods to assess the prediction accuracy and generalization across two datasets. The comparison demonstrates that, on average, existing methods do not predict expression to a high correlation for both HVGs and marker genes. For top-performing methods – Hist2ST, BLEEP, and STimage – the majority of genes are predicted with a correlation between 0.1 and 0.3. We observe slightly higher average performance when predicting the set of marker genes, indicating that useful biological markers may be predicted to a somewhat higher degree of accuracy. Thus, prior selection of useful genes and use of a smaller subset of genes may be beneficial in general to improve performance and reduce computational cost.

We did not observe substantial differences between the overall performance of the model when applying to the Visium and HER2+ datasets, despite the fact that the Visium dataset has an increased resolution and spot density. Although the total number of Visium spots is nearly double that in the HER2+ dataset, the number of independent Visium samples is lower (36 vs 9 patients), possibly reducing the generalization capability as spots from the same sample are correlated [24].

Our assessment of the generalizability and robustness of each method on ID validation and OOD test sets indicate that each method has similar performance on both sets, with the exception of Hist2ST, which shows a drop-off in performance in the OOD set. This may be in part due to the higher model complexity of Hist2ST. However, the relationship between the number of parameters and overfitting is not simple [25], and the performance and generalization may be in part limited by the lack of training data, which only consisted of 4 samples. The other top-performing methods BLEEP and STimage demonstrate robust and consistent performance in the OOD set, which may be due to their different methods of inference that contrast them from the other methods that directly produce point-wise predictions. For STimage, learning the negative binomial distributions for each gene allows the model to account for noise and variance present in the data. For BLEEP, performing inference by querying and using a weighted combination of existing values may be more robust than using a parameterized function, which only indirectly contains information about the true distributions of genes in the training data. These results and observations can provide insight into how deep learning models can be designed to take into account noise and uncertainty in the data and improve robustness.

Our results provide evidence that current methods that transform gene expression in different ways do not have a discernible impact on the performance of the Hist2ST model. Thus, although prior normalization of counts and variance stabilizing transformations such as log-transformation are usually applied to reduce technical variation [26], we find that there are no significant benefits for improving predictive performance for the currently existing models and data. We observe that normalization of the counts with log-transformation results in a smaller spread in the accuracy across predicted genes, which may be beneficial for stabilising predictions.

Although we have tested various methods for improving the state-of-the-art model Hist2ST, such as stain augmentation and normalization, no improvement in the average LOOCV performance or improved generalization to OOD data was observed. This may be due to the complexity of the relationship between stain levels and gene expression, making the effect of color perturbations on the generalization of models unclear and possibly less effective. We did not find improvement by using an auxiliary classification loss, meaning that the use of tissue type information this way did not improve the model at the task of predicting gene expression. Moreover, using pre-trained feature extractors, or simplifying the architecture with a graph attention network, resulted in worse performance compared to the original model. This suggests that models pre-trained on natural image datasets do not transfer well to the domain of histology slide images, and that the global attention layers in the transformer are important for higher performance.

In terms of scalability, we note that the transformer-based methods Hist2ST and HisToGene do not scale well to data with thousands of spots, such as the Visium data, due to the computation of global attention between spots. Although a workaround was implemented in this work, this sacrificed global attention between all the spots in a sample. In contrast, the other methods that treat spots as independent training samples are more scalable as the batch size is independent. Improvements in efficiency and scalability for deep learning methods should be considered for future developments and will be crucial as ST technology develops, providing increases in spot resolution and total spot counts.

## Limitations

Our benchmarking comparison shows that there is still a large gap between the performance of existing models and effectively usable outcomes. However, there are a number of limitations that should be considered when drawing conclusions from our results.

Firstly, the limited size of the spatial transcriptomic datasets used, which was due to limited availability of publicly accessible data, poses a challenge in drawing comprehensive benchmarking conclusions. In this work, the higher resolution Visium data comprised only 9 independent samples, and the HER2+ dataset only 36. Deep learning methods require a large training and diverse training set to avoid overfitting and generalize effectively [24]. Consequently, caution should be exercised when generalizing the findings and extrapolating the performance of these models to datasets of different sizes. In addition, we focused on datasets consisting of breast cancer tissue samples but did not include data from other types of tissues due to the limited amount of training samples from available data. Thus, it may be possible that our results and conclusions are not generalizable to other cancer or tissue types, where the relationship between morphological features and gene expression in the data is different.

In addition, we note that there could be variation in performance due to the set of genes chosen to train and evaluate the models. Although we have focused on a larger set of highly variable genes and a smaller set of cancer markers and found consistent results between methods, this is only a subset of the available expression panel which consists of over 30,000 genes in total for the Visium data. For current methods, it is infeasible to predict all the possible genes, and is computationally expensive to train or re-train models to predict different or larger sets of genes. Therefore, further work on finding a useful biologically relevant set of genes or a set of genes that can be consistently predicted to high accuracy is highly desirable.

## Conclusions and perspectives

While the idea of cost-effective in-silico transcriptomics is appealing, several intricate challenges hinder the potential for an effective and reliable solution. As described by some authors [11], a fundamental challenge lies in the ill-posed nature of the problem. Histology images and spatial transcriptomics data offer complementary views of tissue composition and gene expression patterns. However, expecting image features to predict the expression of all genes is ambitious, and difficult due to the complex, multifaceted nature of gene regulation. Nevertheless, this challenge prompts researchers to identify and prioritize specific gene categories that are biologically relevant for the intended applications. A collection of genes as signatures for cell types or subtypes or those that are associated with the morphological changes and spatial distribution during cancer progression emerge as crucial candidates, as they hold significant diagnostic and therapeutic implications.

However, the joint relationship between gene distribution and tissue image features is still unclear for the majority of genes, and thus there is a large component of uncertainty in both the choice of genes to train on and the model predictions, with many genes not being able to be predicted reliably by current methods. Most existing methods do not have a way of quantifying the uncertainty in the model predictions, which is important for establishing reliability and robustness when testing on new data [27]. STimage is among the first methods to quantify uncertainty prediction for each gene in each spot [10]. More work and improvements in this area and a better understanding of spatial gene expression and tissue relationships will be important to improve these methods and support their use in practice.

Furthermore, the rapidly evolving landscape of spatial transcriptomic methods introduces additional complexities [28]. New technologies generate data that are different in resolution and fidelity to older data. Experimental artifacts, batch effects, and variations within and across samples can confound prediction models, potentially leading to unreliable results. Addressing these challenges requires robust preprocessing techniques and machine-learning algorithms that can properly separate biological signals from noise [29]. In addition, the development of deep learning methods would benefit greatly from having larger amounts of available data across a diverse range of sample types; efforts to promote the open sharing of spatial transcriptomic data would greatly enhance the robustness and reproducibility of experimental outcomes in this field.

In conclusion, predicting gene expression from histology images and spatial transcriptomics data is a formidable challenge that is still largely unsolved by existing methods. Overcoming the challenging nature of the problem by focusing on a subset of predictable genes, dealing with scarce training data, and navigating the complexities introduced by experimental variability are crucial steps toward achieving an effective solution. Success in this endeavour holds the promise of providing cost-effective solutions for digital pathology.

## Data and code availability

The datasets analysed in this study are available here: human HER2-positive breast tumor ST data https://github.com/almaan/her2st, 10x Genomics Visium data and Swarbrick’s Laboratory Visium data https://doi.org/10.48610/4fb74a9. Our code used to produce the data reported here is available at https://github.com/BiomedicalMachineLearning/DeepHis2Exp.

## Supplementary Materials

### Data preprocessing and augmentation

The default preprocessing and augmentation methods for each method are summarized concisely in Table S1. Below, we provide a more detailed overview of the processing steps applied to both gene expression data and image data for each method.

### ST-Net

ST-Net first transforms the total expression in each spot by adding a pseudocount, then applying a logarithmic transformation. Image patches centred on each spot are extracted, with dimensions of 224×224 pixels. During training, image augmentation techniques are utilized, including random rotations and flips at a 50% probability.

### HisToGene

HisToGene first excludes genes expressed in fewer than 1,000 spots across all sections. For each spot, gene expression values are normalized by calculating the UMI count for each gene relative to the total UMI counts across all genes in that spot. This normalized value is then multiplied by 1,000,000 and log-transformed after adding a pseudocount. The image patches are of size 112×112 pixels, equivalent to the diameter of each spot in the HER2+ breast cancer dataset. Image augmentation is applied by random rotation, horizontal flip, and color jitter. *Hist2ST*

Hist2ST follows the same preprocessing of images and gene expression as in HisToGene. Unlike HisToGene, Hist2ST uses data augmentation in training, specifically, in the self-distillation strategy. This is done by creating five randomly perturbed image views of the original image patch. These augmented views are generated using random grayscale, rotation, and horizontal flip transformations.

### STimage

STimage preprocesses the gene counts by log transformation after adding a pseudocount. Image patches of size 299×299 pixels are extracted from each spot. Stain normalization using the Vahadane method [30] is conducted on the tiles to ensure colour distribution patterns are akin to a template image. Tiles with low tissue coverage (less than 70%) are removed. For training, image augmentation techniques are utilized, including random flipping, scaling, rotation, blurring, adding noise, and colour jitter.

### DeepSpaCE

DeepSpaCE preprocesses the data by filtering out spots with low UMI counts or a low number of measured genes. UMI counts are normalized using the SCTransform [31] function from the Seurat package, and expression values are adjusted through min-max scaling. Images of size 224×224 pixels are used, and those with high mean RGB values, above 80%, are excluded. Image augmentation is performed using diverse transformations, including flipping, cropping, noise addition, blurring, distortion, contrast adjustment, and colour-shifting, to enhance the model’s performance and adaptability.

### BLEEP

The BLEEP workflow involves normalizing the expression levels of each spot by dividing them by the total count and applying a log transformation. To address batch effects, the Harmony algorithm is utilized to adjust the expressions of the samples. Image patches with dimensions of 224×224 pixels are extracted around each spot, and during training, they are augmented by random flips and rotations to enhance the model’s performance.

**Table S1.** Summary of data preprocessing methods from six methods.

| Model | Tile size (pixels) | Data augmentation | Gene expression Pre-processing |
| --- | --- | --- | --- |
| ST-Net | 224×224 | Random Rotation & Flipping | Log Transformation |
| HisToGene | 112×112 | Random Rotation & Flipping + ColorJitter | Normalization + Log Transformation |
| Hist2ST | 112×112 | Random Rotation & Flipping + ColorJitter (Self-distillation strategy) | Normalization + Log Transformation |
| STimage | 299×299 | Color Normalization (Vahadane) + Random one of flipping, cropping, noise addition, blurring, distortion, contrast adjustment, colour-shifting + Remove tiles with low tissue coverage (< 70%) | Log Transformation |
| DeepSpaCE | 224×224 | Remove tiles with high RGB values | SCTransform + MinMax Scaling |
| BLEEP | 224×224 | Random rotation & flip | Normalization + Log Transformation |

**Table S2.** Summary of image encoders and loss functions from six models.

| Model | Image Encoder | Loss Function |
| --- | --- | --- |
| ST-Net | Densenet121 (Pre-trained) | Mean Squared Error |
| HisToGene | MLP + Transformer | Mean Squared Error |
| Hist2ST | CNN + Transformer + GCN | Mean Squared Error (Regression) + Mean Squared Error (Self-distillation) + Negative log likelihood |
| STimage | Resnet50 (Pre-trained) | Negative log likelihood |
| DeepSpaCE | VGG16 (Pre-trained) | Mean Squared Error |
| BLEEP | Resnet50 (Pre-trained) | Cross Entropy |

**Table S3.** List of breast cancer datasets.

| No. | Name | URL | Technology |
| --- | --- | --- | --- |
| BC0 | Human breast cancer in situ capturing transcriptomics | https://tinyurl.com/355myrts | Legacy ST |
| BC1 | Human HER2-positive breast tumor ST data | https://github.com/almaan/her2st | Legacy ST |
| BC2 | 10x Genomics breast cancer dataset | https://tinyurl.com/ywtfbctr | Visium |
| BC3 | Swarbrick breast cancer dataset | https://tinyurl.com/2xboavb3 | Visium |

**Table S4.**
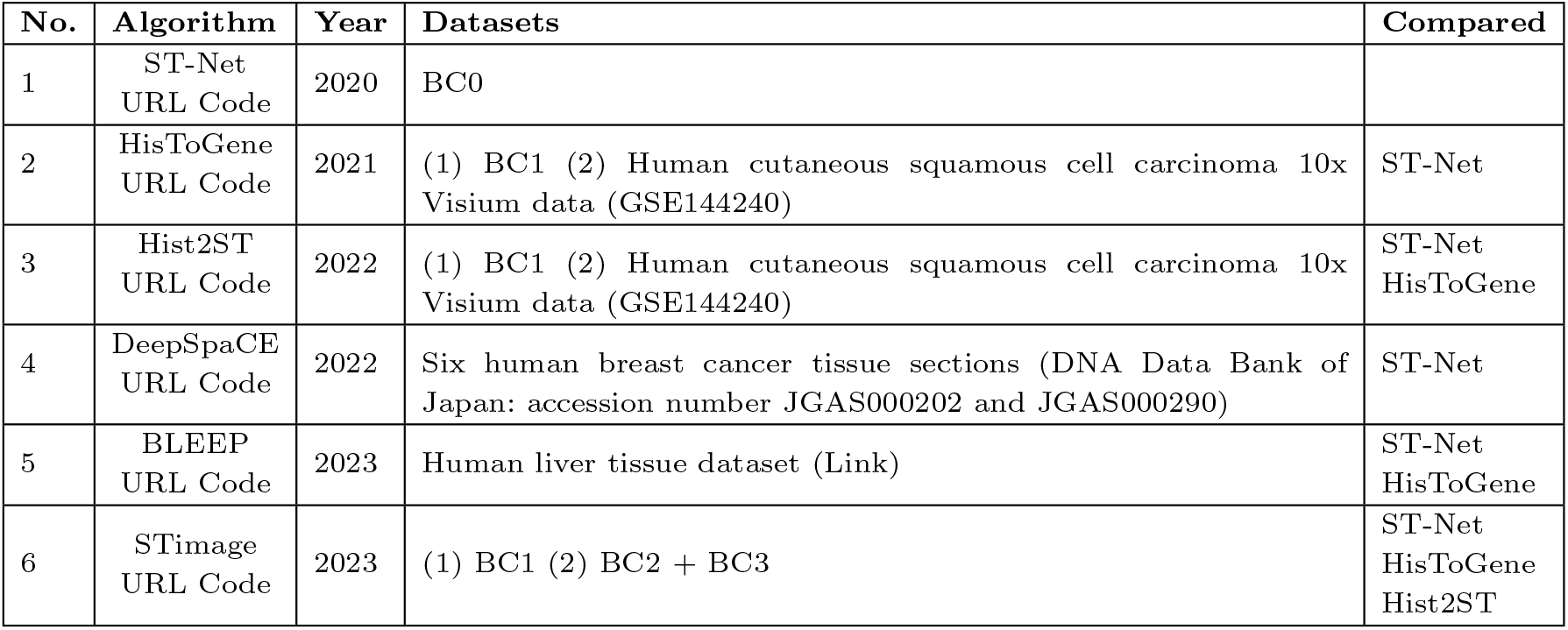
Published algorithms and datasets. See Table S3 for abbreviated breast cancer dataset details.

| No. | Algorithm | Year | Datasets | Compared |
| --- | --- | --- | --- | --- |
| 1 | ST-Net<br>URL Code | 2020 | BC0 |  |
| 2 | HisToGene<br>URL Code | 2021 | (1) BC1 (2) Human cutaneous squamous cell carcinoma 10x Visium data (GSE144240) | ST-Net |
| 3 | Hist2ST<br>URL Code | 2022 | (1) BC1 (2) Human cutaneous squamous cell carcinoma 10x Visium data (GSE144240) | ST-Net<br>HisToGene |
| 4 | DeepSpaCE<br>URL Code | 2022 | Six human breast cancer tissue sections (DNA Data Bank of Japan: accession number JGAS000202 and JGAS000290) | ST-Net |
| 5 | BLEEP<br>URL Code | 2023 | Human liver tissue dataset (Link) | ST-Net<br>HisToGene |
| 6 | STimage<br>URL Code | 2023 | (1) BC1 (2) BC2 + BC3 | ST-Net<br>HisToGene<br>Hist2ST |

**Fig. S1.**
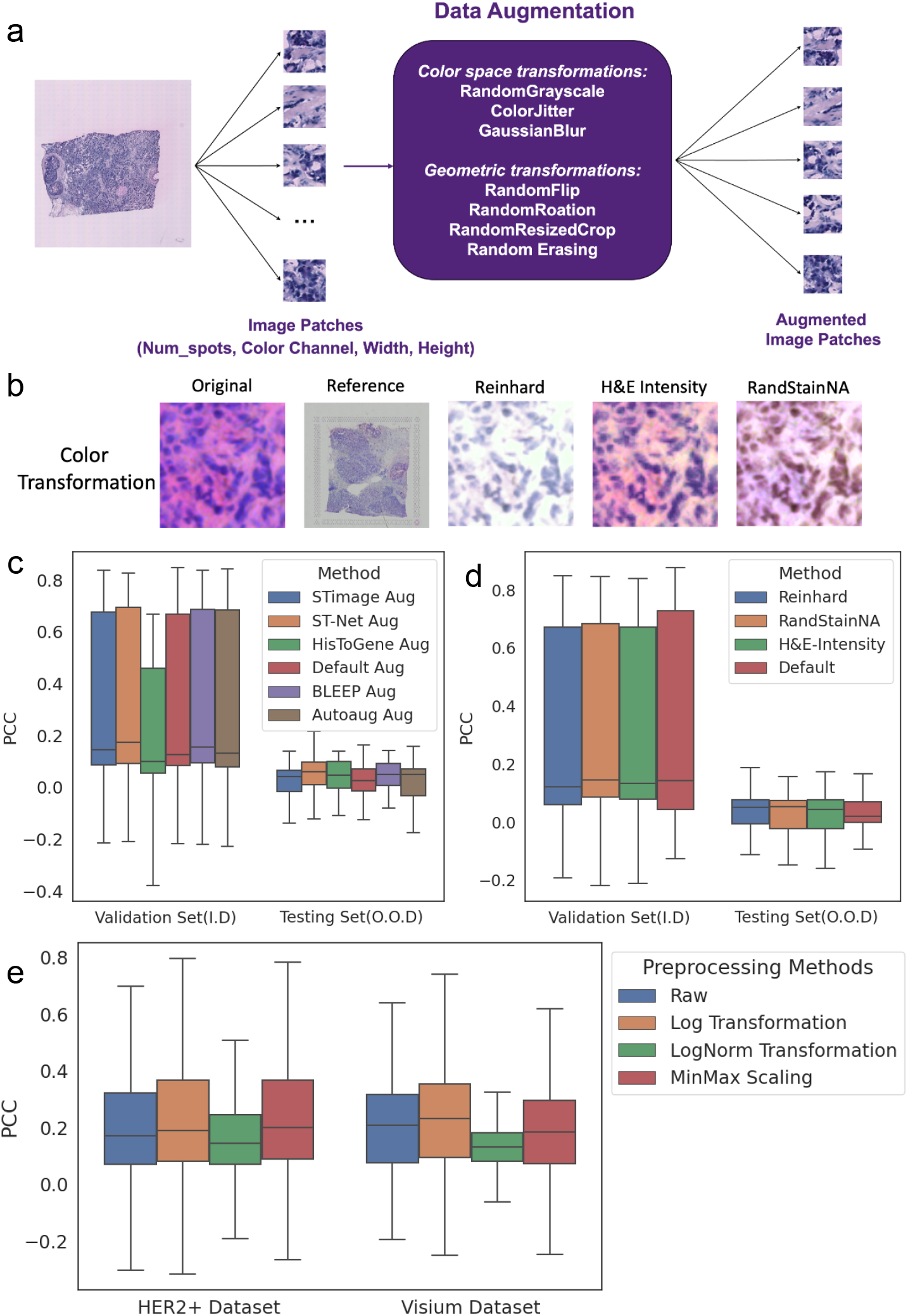
a) Summary of image augmentation techniques used by different methods. **b)** Demonstration of different colour augmentation and stain normalization methods, for example, H&E image patch. **c-e)** Comparing augmentation and gene expression processing techniques using Hist2ST as the base model. **c)** Evaluation of augmentation methods on the generalization ability of the Hist2ST model using in-distribution validation set (Swarbrick data samples) and out-of-distribution data (10x Genomics data samples). **d)** Evaluation of color augmentation and normalization methods on the generalization ability of the Hist2ST model (validation and testing set same as in c). **e)** Evaluation of the effect of gene expression preprocessing on Hist2ST model performance by LOOCV. Box plots show the distribution of PCC values for the 12 marker genes.

**Fig. S2.**
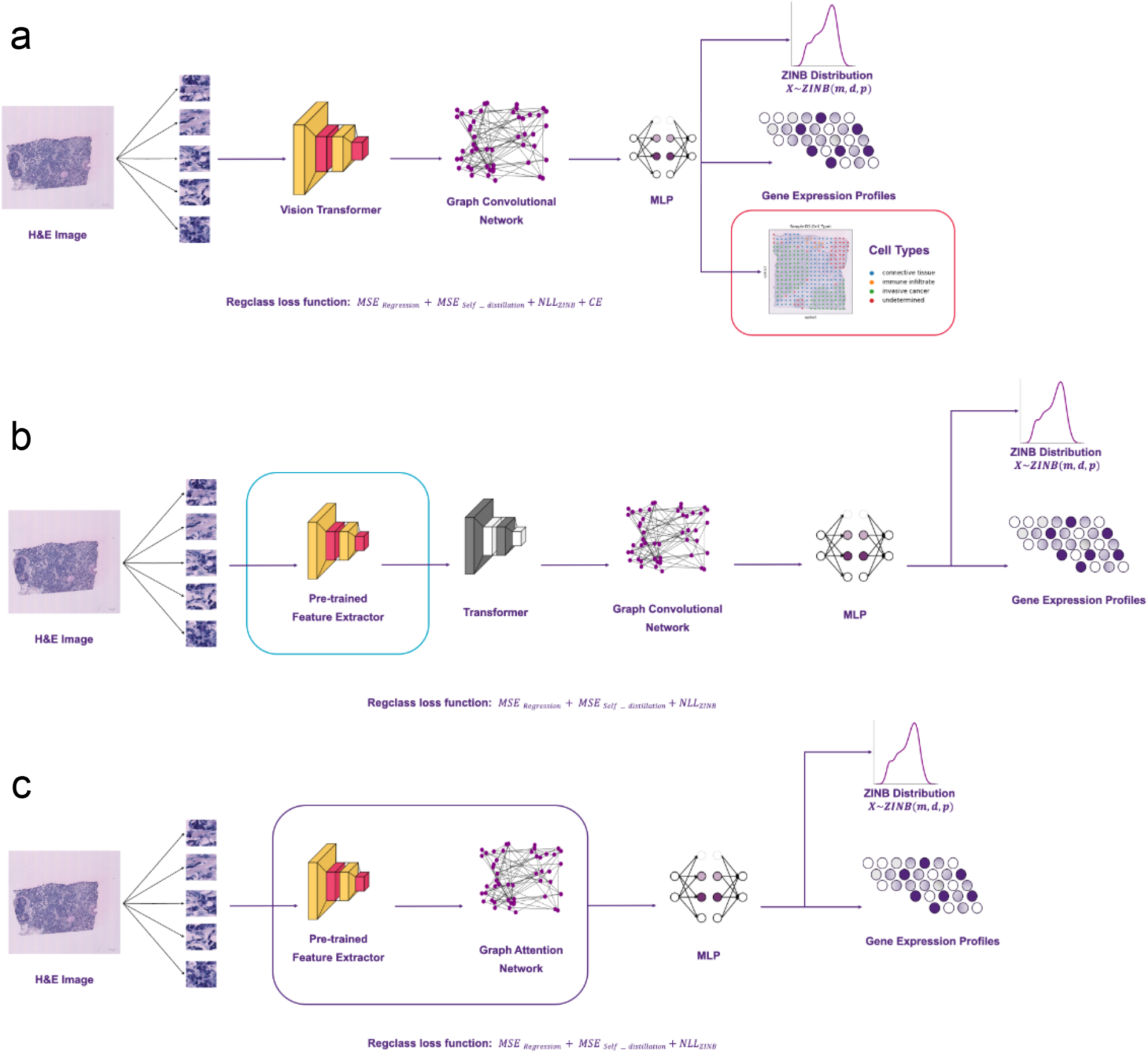
Workflow of Hist2ST variants. **a)** The Hist2ST model with auxiliary classification loss (Regclass). The red frame highlights the addition of the classifier module. The extracted features are shared for the cell type classification head and gene expression prediction head. **b)** The architecture of the Hist2ST model with pre-trained image encoders. The blue frame highlights the modification, replacing the Convmixer module with pre-trained models (Swin-Transformer, ResNet-50, EfficientNetV2) model. **c)** Simplified Hist2ST architecture based on previous pre-trained feature extractor modification, and further replacing the Transformer and Graph Convolutional Network with the Graph Attention Network (purple frame).

**Fig. S3.**
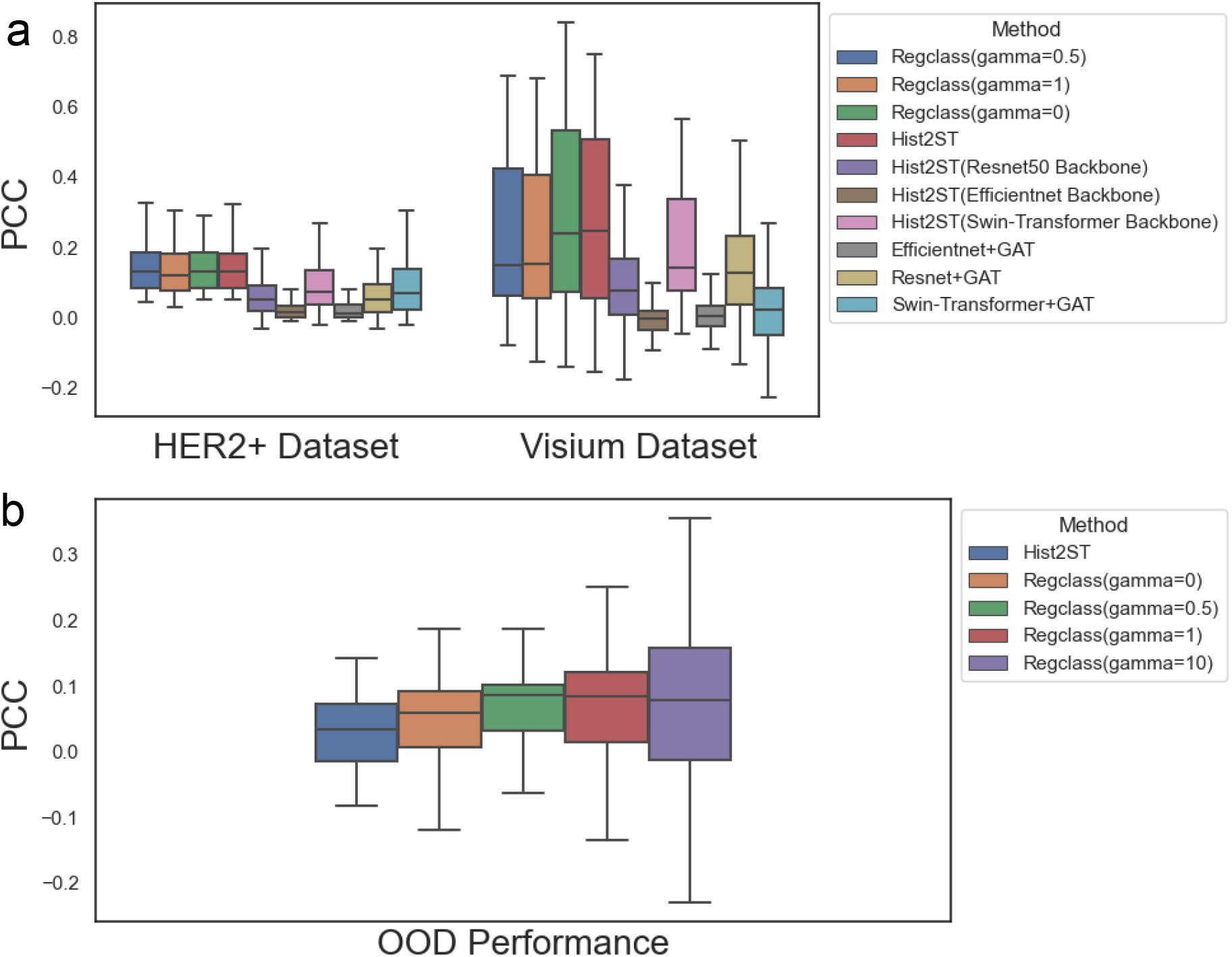
a) Performance comparison of Hist2ST variants on the HER2+ and Visium dataset. The variants include Regclass, Hist2ST with 3 different pre-trained feature extractors (Resnet50, Efficientnet v2, Swin-transformer) and Hist2ST with combined pre-trained feature extractors and graph attention network. **b)** Generalization performance of Regclass on the out-of-distribution dataset, with different values of the gamma hyper-parameter. Box plots show the distribution of PCC values for the 12 marker genes.

**Fig. S4.**
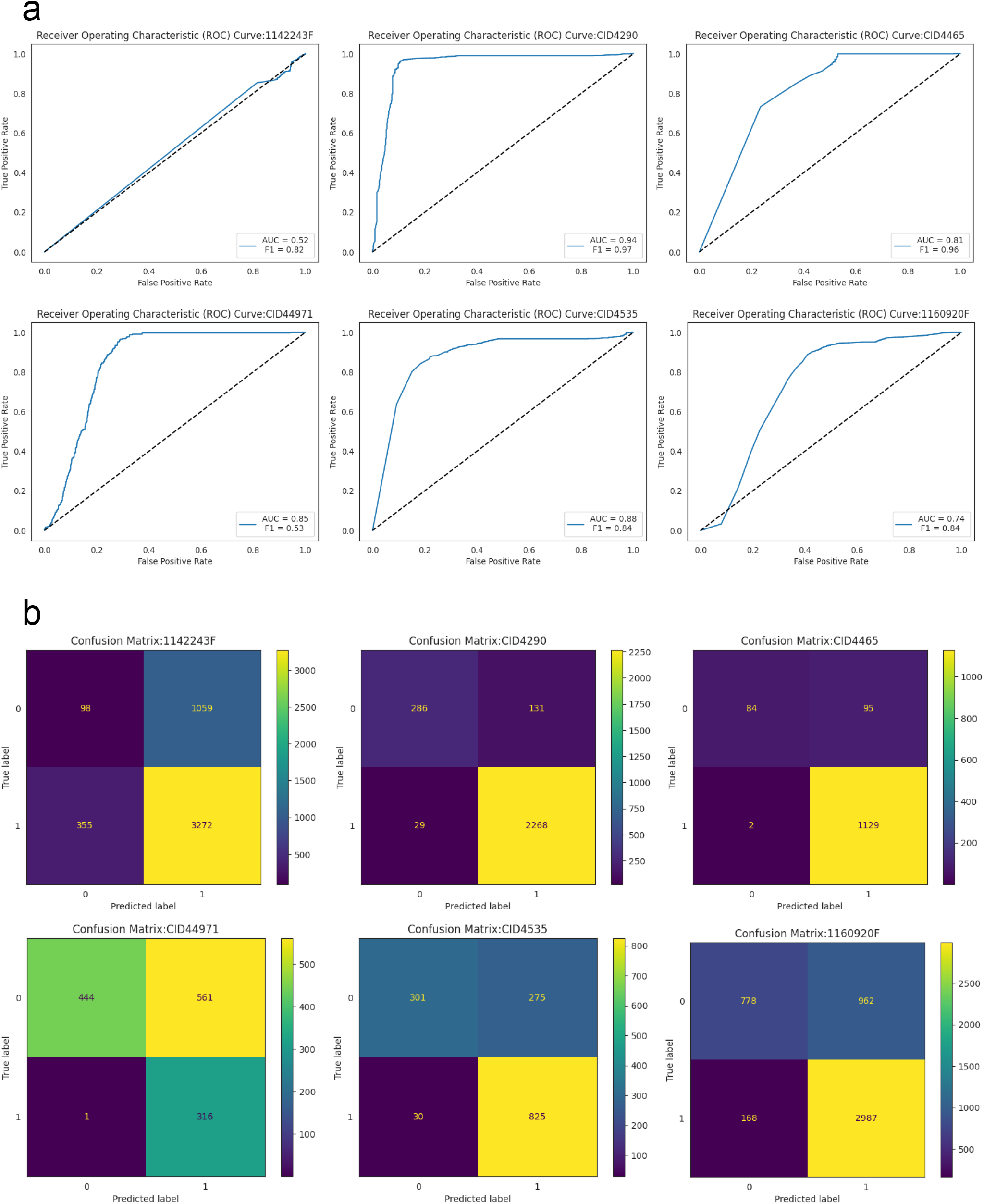
Classification performance of Regclass on Visium dataset (6 samples with annotations) under leave-one-out cross-validation. **a)** Receiver operating characteristic curve and F1 score for each sample. **b)** Confusion matrix for each sample.

## References

1. Vivien Marx. Method of the Year: spatially resolved transcriptomics. 2021. Nature Methods, 18(1):9–14, January 2021.

2. Patrik L. Ståhl Fredrik Salmén, Sanja Vickovic, Anna Lundmark, José Fernández Navarro, Jens Magnusson, Stefania Giacomello, Michaela Asp, Jakub O. Westholm, Mikael Huss, Annelie Mollbrink, Sten Linnarsson, Simone Codeluppi, åke Borg, Fredrik Pontén, Paul Igor Costea, Pelin Sahlén, Jan Mulder, Olaf Bergmann, Joakim Lundeberg, and Jonas Frisén. Visualization and analysis of gene expression in tissue sections by spatial transcriptomics. Science (New York, N.Y.), 353(6294):78–82, July 2016.

3. Qichao Yu, Miaomiao Jiang, and Liang Wu. Spatial transcriptomics technology in cancer research. Frontiers in Oncology, page 5486, 2022.

4. Benoît Schmauch, Alberto Romagnoni, Elodie Pronier, Charlie Saillard, Pascale Maillé, Julien Calderaro, Aurélie Kamoun, Meriem Sefta, Sylvain Toldo, Mikhail Zaslavskiy, Thomas Clozel, Matahi Moarii, Pierre Courtiol, and Gilles Wainrib. A deep learning model to predict RNA-Seq expression of tumours from whole slide images. Nature Communications, 11(1):3877, August 2020.

5. Yinxi Wang, Kimmo Kartasalo, Philippe Weitz, Balázs Ács, Masi Valkonen, Christer Larsson, Pekka Ruusuvuori, Johan Hartman, and Mattias Rantalainen. Predicting Molecular Phenotypes from Histopathology Images: A Transcriptome-Wide Expression-Morphology Analysis in Breast Cancer. Cancer Research, 81(19):5115–5126, October 2021.

6. Bryan He, Ludvig Bergenstråhle, Linnea Stenbeck, Abubakar Abid, Alma Andersson, åke Borg, Jonas Maaskola, Joakim Lundeberg, and James Zou. Integrating spatial gene expression and breast tumour morphology via deep learning. Nature biomedical engineering, 4(8):827–834, 2020.

7. Minxing Pang, Kenong Su, and Mingyao Li. Leveraging information in spatial transcriptomics to predict superresolution gene expression from histology images in tumors. bioRxiv, pages 2021–11, 2021.

8. Yuansong Zeng, Zhuoyi Wei, Weijiang Yu, Rui Yin, Yuchen Yuan, Bingling Li, Zhonghui Tang, Yutong Lu, and Yuedong Yang. Spatial transcriptomics prediction from histology jointly through transformer and graph neural networks. Briefings in Bioinformatics, 23(5):bbac297, 2022.

9. Taku Monjo, Masaru Koido, Satoi Nagasawa, Yutaka Suzuki, and Yoichiro Kamatani. Efficient prediction of a spatial transcriptomics profile better characterizes breast cancer tissue sections without costly experimentation. Scientific Reports, 12(1):4133, 2022.

10. Xiao Tan, Onkar Mulay, Samual MacDonald, Taehyun Kim, Jason Werry, Peter T Simpson, Fred Roosta, Maciej Trzaskowski, and Quan Nguyen. Stimage: robust, confident and interpretable models for predicting gene markers from cancer histopathological images. bioRxiv, pages 2023–05, 2023.

11. Ronald Xie, Kuan Pang, Gary D Bader, and Bo Wang. Spatially resolved gene expression prediction from h&e histology images via bi-modal contrastive learning. arXiv preprint arXiv:2306.01859, 2023.

12. Kaiming He, Xiangyu Zhang, Shaoqing Ren, and Jian Sun. Deep residual learning for image recognition. In Proceedings of the IEEE conference on computer vision and pattern recognition, pages 770–778, 2016.

13. Jia Deng, Wei Dong, Richard Socher, Li-Jia Li, Kai Li, and Li Fei-Fei. Imagenet: A large-scale hierarchical image database. In 2009 IEEE conference on computer vision and pattern recognition, pages 248–255. Ieee, 2009.

14. Gökcen Eraslan, Lukas M Simon, Maria Mircea, Nikola S Mueller, and Fabian J Theis. Single-cell rna-seq denoising using a deep count autoencoder. Nature communications, 10(1):390, 2019.

15. Yixiao Ge, Xiao Zhang, Ching Lam Choi, Ka Chun Cheung, Peipei Zhao, Feng Zhu, Xiaogang Wang, Rui Zhao, and Hongsheng Li. Self-distillation with batch knowledge ensembling improves imagenet classification. arXiv preprint arXiv:2104.13298, 2021.

16. Ting Chen, Simon Kornblith, Mohammad Norouzi, and Geoffrey Hinton. A simple framework for contrastive learning of visual representations. In International conference on machine learning, pages 1597–1607. PMLR, 2020.

17. Alma Andersson, Ludvig Larsson, Linnea Stenbeck, Fredrik Salmén, Anna Ehinger, Sunny Z Wu, Ghamdan Al-Eryani, Daniel Roden, Alex Swarbrick, åke Borg, et al. Spatial deconvolution of her2-positive breast cancer delineates tumor-associated cell type interactions. Nature communications, 12(1):6012, 2021.

18. Amanda Janesick, Robert Shelansky, Andrew D. Gottscho, Florian Wagner, Morgane Rouault, Ghezal Beliakoff, Michelli Faria de Oliveira, Andrew Kohlway, Jawad Abousoud, Carolyn A. Morrison, Tingsheng Yu Drennon, Seayar H. Mohabbat, Stephen R. Williams, 10x Development Teams, and Sarah E. B. Taylor. High resolution mapping of the breast cancer tumor microenvironment using integrated single cell, spatial and in situ analysis of FFPE tissue, November 2022.

19. Sunny Z. Wu, Ghamdan Al-Eryani, Daniel Roden, Simon Junankar, Kate Harvey, Alma Andersson, Aatish Thennavan, Chenfei Wang, James Torpy, Nenad Bartonicek, Taopeng Wang, Ludvig Larsson, Dominik Kaczorowski, Neil I. Weisenfeld, Cedric R. Uytingco, Jennifer G. Chew, Zachary W. Bent, Chia-Ling Chan, Vikkitharan Gnanasambandapillai, Charles-Antoine Dutertre, Laurence Gluch, Mun N. Hui, Jane Beith, Andrew Parker, Elizabeth Robbins, Davendra Segara, Caroline Cooper, Cindy Mak, Belinda Chan, Sanjay Warrier, Florent Ginhoux, Ewan Millar, Joseph E. Powell, Stephen R. Williams, X. Shirley Liu, Sandra O’Toole, Elgene Lim, Joakim Lundeberg, Charles M. Perou, and Alexander Swarbrick. A single-cell and spatially resolved atlas of human breast cancers. Nature genetics, 53(9):1334–1347, September 2021.

20. Sebastian Otálora, Niccoló Marini, Damian Podareanu, Ruben Hekster, David Tellez, Jeroen Van Der Laak, Henning Müller, and Manfredo Atzori. Stainlib: a python library for augmentation and normalization of histopathology h&e images. bioRxiv, pages 2022–05, 2022.

21. Yiqing Shen, Yulin Luo, Dinggang Shen, and Jing Ke. Randstainna: Learning stain-agnostic features from histology slides by bridging stain augmentation and normalization. In International Conference on Medical Image Computing and Computer-Assisted Intervention, pages 212–221. Springer, 2022.

22. Mingxing Tan and Quoc Le. Efficientnet: Rethinking model scaling for convolutional neural networks. In International conference on machine learning, pages 6105–6114. PMLR, 2019.

23. Ze Liu, Yutong Lin, Yue Cao, Han Hu, Yixuan Wei, Zheng Zhang, Stephen Lin, and Baining Guo. Swin transformer: Hierarchical vision transformer using shifted windows. In Proceedings of the IEEE/CVF international conference on computer vision, pages 10012–10022, 2021.

24. Kenji Kawaguchi, Leslie Pack Kaelbling, and Yoshua Bengio. Generalization in Deep Learning. pages 112–148. December 2022. arXiv:1710.05468 [cs, stat].

25. Chiyuan Zhang, Samy Bengio, Moritz Hardt, Benjamin Recht, and Oriol Vinyals. Understanding deep learning requires rethinking generalization. CoRR, abs/1611.03530, 2016.

26. Yingdong Zhao, Ming-Chung Li, Mariam M. Konaté, Li Chen, Biswajit Das, Chris Karlovich, P. Mickey Williams, Yvonne A. Evrard, James H. Doroshow, and Lisa M. McShane. Tpm, fpkm, or normalized counts? A comparative study of quantification measures for the analysis of rna-seq data from the nci patient-derived models repository. Journal of Translational Medicine, 19(1):269, June 2021.

27. Moloud Abdar, Farhad Pourpanah, Sadiq Hussain, Dana Rezazadegan, Li Liu, Mohammad Ghavamzadeh, Paul Fieguth, Xiaochun Cao, Abbas Khosravi, U. Rajendra Acharya, Vladimir Makarenkov, and Saeid Nahavandi. A review of uncertainty quantification in deep learning: Techniques, applications and challenges. Information Fusion, 76:243–297, December 2021.

28. Jun Du, Yu-Chen Yang, Zhi-Jie An, Ming-Hui Zhang, Xue-Hang Fu, Zou-Fang Huang, Ye Yuan, and Jian Hou. Advances in spatial transcriptomics and related data analysis strategies. Journal of Translational Medicine, 21(1):330, May 2023.

29. Shuangsang Fang, Bichao Chen, Yong Zhang, Haixi Sun, Longqi Liu, Shiping Liu, Yuxiang Li, and Xun Xu. Computational Approaches and Challenges in Spatial Transcriptomics. Genomics, Proteomics & Bioinformatics, 21(1):24–47, February 2023.

30. Abhishek Vahadane, Tingying Peng, Amit Sethi, Shadi Albarqouni, Lichao Wang, Maximilian Baust, Katja Steiger, Anna Melissa Schlitter, Irene Esposito, and Nassir Navab. Structure-preserving color normalization and sparse stain separation for histological images. IEEE transactions on medical imaging, 35(8):1962–1971, 2016.

31. Christoph Hafemeister and Rahul Satija. Normalization and variance stabilization of single-cell rna-seq data using regularized negative binomial regression. Genome biology, 20(1):296, 2019.

